# Mineral deposition and vascular invasion of hydroxyapatite reinforced collagen scaffolds seeded with human adipose-derived stem cells

**DOI:** 10.1101/691618

**Authors:** Holly E. Weiss-Bilka, Matthew J. Meagher, Joshua A. Gargac, Glen L. Niebur, Ryan K. Roeder, Diane R. Wagner

## Abstract

Collagen-based scaffolds reinforced with hydroxyapatite (HA) are an attractive choice for bone tissue engineering because their composition mimics that of bone. We previously reported the development of compression-molded collagen-HA scaffolds that exhibited high porosity, interconnected pores, and mechanical properties that were well-suited for surgical handling and fixation. The objective of this study was to investigate the use of these novel collagen-HA scaffolds in combination with human adipose-derived stem cells (hASCs) as a template for bone formation in a subcutaneous athymic mouse model. Cell-seeded constructs were pre-treated with either control or osteogenic media. Cell-free and collagen-only groups were included, as was a clinically approved bone void filler as a control for the material. After 8 weeks implantation, cell-free collagen-HA scaffolds and those that were pre-seeded with osteogenically differentiated hASCs supported bone formation and vascular invasion at comparable rates. HA-reinforcement allowed collagen constructs to maintain their implanted shape, provided for improved cell-tissue-scaffold integration, and resulted in a more organized tissue when pre-treated in an osteogenic induction medium. Scaffold type and pre-treatment also determined osteoclast activity and therefore potential remodeling of the constructs. Results suggest that it may be necessary to match the scaffold with a particular cell type and cell-specific pre-treatment to achieve optimal bone formation.

## Introduction

Non-union fractures and critical size bone defects have a considerable impact upon the global population [1]. In fact, bone is the second most transplanted tissue worldwide with an estimated 2.2 million graft procedures performed each year [2]. Yet despite their prevalence, autogenic grafts are limited by the availability of donor tissue and are often associated with donor site pain, while allogenic grafting techniques carry the risk of morbidity and infection [3]. These limitations have inspired many research efforts involving laboratory-produced tissue replacements; however current approaches to bone tissue engineering generally lack a sufficient functionality compared to native bone matrix.

Collagen-based scaffolds reinforced with hydroxyapatite (HA) are an attractive choice for bone tissue engineering because they mimic the key components of bone, collagen and mineral, and possess improved mechanical properties compared to either component alone [4,5]. We previously reported novel compression-molded collagen-HA scaffolds that exhibited high porosity (85 – 90%), ∼300 – 400 μm interconnected pores, struts composed of high-density collagen fibrils reinforced with HA whiskers, and mechanical properties that were well-suited for surgical handling and fixation [6]. These scaffolds were also conducive to the infiltration and *in vitro* differentiation of adipose-derived stem cells [6]. After ectopic implantation. the vascular density, cell density, matrix deposition, and micro-CT bone volume increased with increasing HA content in the scaffolds [7]. However, in these previous studies, the collagen-HA scaffolds were implanted without pre-seeding with osteogenic cells, which may further increase bone tissue generation.

Human adult adipose-derived stem cells (hASCs) are an appealing complement to such scaffolds because they are abundant and have been shown to contribute to both bone formation and vasculogenesis [8] *in vivo*. Subcutaneous implantation in immunodeficient mice is an established model for evaluating combinations of hASCs with various scaffolds and culture conditions, with multiple reports of successful bone formation in the literature [9]. One of the earliest successful studies coupled β-tricalcium phosphate discs with hASCs and then pretreatment in an osteoinductive medium for 2 weeks. During 8 weeks of subcutaneous implantation in nude mice, the cell-seeded discs developed an osteocalcin-rich tissue containing osteoclasts and infiltrated by blood vessels [10]. Another study reported bone formation in 4 of 5 HA-tricalcium phosphate (TCP) and in 1 of 5 Collagraft^®^ (collagen-HA-TCP composite matrix) scaffolds that had been seeded with untreated hASCs and subjected to 6 weeks subcutaneous implantation in nude mice [11]. In a different investigation, hASCs seeded onto porous HA ceramic scaffolds and cultured in a 3-D perfusion system for 5 days prior to subcutaneous implantation in nude mice for 8 weeks resulted in well vascularized constructs containing osteoprogenitor cells and positive immunostaining for human bone sialoprotein [12]. Interestingly, immunostaining for human CD31 and CD34 indicated that newly formed vessels were of human origin. More recently, the importance of surface topography was demonstrated in the context of TiO_2_ nanotube surfaces, which enhanced the osteogenic differentiation of hASCs both *in vitro* and *in vivo* [13]. Finally, hASCs were first cultured on either extracellular matrix derived from bone marrow-derived mesenchymal stem cells or tissue culture plastic before being loaded on HA powder and implanted subcutaneously in immunodeficient mice; hASCs that had been expanded on the cell-derived matrix produced more bone tissue compared to those cultured on tissue culture plastic [14].

The objective of this study was therefore to investigate the use of novel collagen-HA (CHA) scaffolds in combination with hASCs as a template for bone formation in a subcutaneous athymic mouse model. Cell-seeded constructs were pre-treated with either control medium (CM), or an osteogenic medium (OM). A cell-free, or acelluar (Acel), control group of collagen-HA scaffolds was included to assess the osteoinductive capacity of the scaffold itself. This group was cultured in OM for the same duration as cell-seeded groups. The effect of HA in the scaffold was examined by including a collagen-only (Col) control group. Finally, NuOss™ (Nu) – a clinically approved bone void filler previous reported to support bone formation in ectopic models when coupled with human periosteum-derived cells, human mesoangioblasts and a murine pre-chondrogenic cell line [15,16] – was included as a control for the scaffold material.

## Methods

### Cell culture

Human adipose-derived stem cells (ZenBio, Durham, NC) derived and pooled from the subcutaneous adipose tissue of 5 non-diabetic female donors were expanded as previously described [17,18]. During expansion, cells were plated at a density of 3,000 cells/cm^2^ and were maintained in DMEM/F12 medium (MediaTech, Herndon, VA) containing 10 % FBS (Atlas Biologicals, Fort Collins, CO), 1 % Penicillin-Streptomycin (Pen-Strep, MediaTech), 5 ng/mL human epidermal growth factor (hEGF), 1 ng/mL human fiberblast growth factor-2 (hFGF2) and 0.25 ng/mL transforming growth factor-β1 (TGF-β1; all growth factors from PeproTech, Rocky Hill, NJ). Cells were treated with fresh medium every two days.

### Scaffold preparation

Collagen (Col) and collagen with 40 vol % hydroxyapatite whiskers (CHA) scaffolds with 85 % porosity and a mean pore size of ∼375 µm were fabricated as previously described [6,7]. NuOss™ scaffolds were purchased from Ace Surgical Supply (Brockton, MA). All scaffolds were sized to 21 mm^3^ (3 mm diameter × 3 mm height) using a sterile biopsy punch. After crosslinking and porogen leaching of Col and CHA scaffolds, implants were sterilized by submersion in 70 % ethanol and rehydrated in sterile PBS followed by sterile culture medium supplemented with 1 % Pen-Strep.

### Cell seeding and *in vitro* culture

For cell seeding, pre-sterilized scaffolds were transferred to a sterile gauze pad, where excess medium was removed, and then to sterile, agarose-coated tissue culture plates. Passage 7 hASCs were drop-seeded onto each scaffold in a 20 µL suspension at a seeding density of 21 × 10^6^ cells/mL and placed into a humidified incubator at 37 °C and 5 % CO_2_ for one hour to attach. After one hour, 1 mL of either control medium (*CM*: DMEM-HG, 10 % FBS, 1 % Pen-Strep) or osteogenic differentiation medium [10,19] (*OM*: CM supplemented with 50 µg/mL ascorbic acid, 10 mM β–glycerophosphate, 0.1 µM dexamethasone; all additives from Sigma-Aldrich, St. Louis, MO) was added to each construct. NuOss™ scaffolds, previously successful in inducing bone formation with human periosteal-derived cells in an ectopic model [15], were included as a control. These scaffolds are of bovine origin and contain natural bone mineral in an open collagen network. Preliminary studies demonstrated that no bone was formed when this scaffold was implanted without cells. Collagen-only (Col) scaffolds were included to assess the contribution of HA whiskers to bone formation, and the importance of pre-implantation osteogenic induction was investigated by treating seeded hASCs with OM compared to CM.

Following overnight incubation, the culture medium was collected from each well containing a cell-seeded scaffold and unattached cells were pelleted at 300 × *g*, re-suspended, and counted with a hemocytometer to determine seeding efficiency. Fresh culture medium was added to all scaffolds three times per week for 14 days, which is the time at which hASC monolayer cultures have previously shown markers of osteogenic differentiation *in vitro* [11,19]. ALP activity and calcium nodules were confirmed in cells grown in monolayer and treated with OM and CM for 14 days. On the day of implantation, 1 scaffold per group (see Table 1) was rinsed with PBS, fixed overnight in 4 % paraformaldehyde at 4 °C, embedded in optimal cutting temperature (OCT) compound and processed for histology.

**Table 1.** Overview of experimental groups.

| *Abbreviation* | Nu OM | Col OM | CHA Acel | CHA CM | CHA OM |
| --- | --- | --- | --- | --- | --- |
| *Scaffold* | NuOss™ | Collagen | Collagen/HA | Collagen/HA | Collagen/HA |
| *Calcium source* | CaP granules | - | HA whiskers | HA whiskers | HA whiskers |
| *hASCs* | + | + | - | + | + |
| *Medium* | OM | OM | OM | CM | OM |

### Subcutaneous ectopic implantation in mice

Remaining scaffolds were implanted subcutaneously in the cervical region of athymic nude mice (Harlan Laboratories, Indianapolis, IN). All procedures were carried out in compliance with protocols approved by the Institutional Animal Care and Use Committee (IACUC) of the University of Notre Dame. Mice were anesthetized with a ‘rodent cocktail’ consisting of 100 mg/mL Ketamine, 20 mg/mL Xylazine and 10 mg/mL Acepromazine (all from Henry Schein, Dublin, OH) in sterile saline, according to the following dosage: Volume anesthetic [µL] = (Body weight [g] × 10) – 50. Each mouse received three constructs through a small incision in the center of the dorsal region. Following 8 weeks of implantation, mice were sacrificed and scaffolds were recovered.

### Micro-computed tomography of scaffolds and explants

Prior to cell seeding, all scaffolds to be implanted were scanned by micro-computed tomography (micro-CT; Bioscan NanoSPECT/CT, Mediso Medical Imaging Systems, Budapest, Hungary) at 10 µm resolution, 70 kVp voltage, 100 mA current with 720 projections at 600 ms integration time. Micro-CT images were median filtered to reduce noise. The bone volume (BV) was measured by segmenting images at a threshold of 1900, which corresponded to 294 mg HA/cm^3^ using a custom calibration phantom [20]. An *ex-vivo* micro-CT scan was performed on all explants after overnight fixation in 4 % paraformaldehyde, using the same parameters as the pre-implantation scans. De novo bone formation was measured by the percent difference in thresholded BV between the implant and explant.

### Histology and immunofluorescence

Fixed samples were rinsed with PBS, decalcified in a 0.5 M EDTA solution and subjected to a series of increasing concentrations of sucrose in PBS. Explants were equilibrated in OCT compound (Sakura, Torrance, CA) for 3 hours, frozen in dry ice-cooled isopentane, and stored at −80 °C prior to sectioning. Each embedded sample was cryosectioned at a thickness of 7 – 9 µm and transferred to gelatin subbed slides, which were dried at 37 °C for 2 hours and then stored at −80 °C. Slides were warmed to room temperature and dried prior to all staining procedures. Sections were stained with H&E following standard histological techniques. Tartrate resistant acid phosphatase (TRAP) staining was performed by 2 hour incubation in TRAP buffer (50 mM sodium acetate, 30 mM sodium tartrate, 0.1 % Triton X-100, pH 5). The buffer was then replaced with TRAP stain for 1 hour before 2 rinses in PBS, counterstaining with hematoxylin and mounting with aqueous mounting medium [16].

All immunofluorescence (IF) procedures were optimized for the specific antibody. Unless otherwise noted, IF procedures were performed for CD31 (R&D Systems; 10 µg/mL), osteopontin (goat polyclonal, R&D Systems; 15 µg/mL), vascular endothelial growth factor (VEGF rabbit polyclonal, Abcam; 1 µg/mL), osteocalcin (OCN rabbit polyclonal, Abcam; 1:1000) as follows. Slides were brought to room temperature, hydrated in PBS and, when necessary, subjected to antigen retrieval consisting of 5 minute incubation at 90 °C in 1 mM EDTA, 0.05% Polysorbate-20, pH 8.0. Sections were cooled to room temperature, rinsed with PBS, and blocked with 1 % normal serum from the animal in which the secondary antibody was raised and 0.3 M glycine in PBS for 30 minutes at room temperature. Primary antibody was applied in blocking solution overnight at 4 °C. Slides were rinsed with PBS before the secondary antibody (Alexa Fluor^®^ anti-goat IgG and anti-rabbit IgG, Invitrogen; 1:200) was diluted in 1 % BSA and applied for 1 hour at room temperature. Finally, sections were counterstained with 4’,6-Diamidino-2-Phenylindole (DAPI), rinsed with PBS, and mounted in an aqueous medium.

For human nuclei antibody (HNA), sections were brought to room temperature, permeabilized with ice cold acetone for 10 minutes at room temperature, dried and re-hydrated in PBS. Blocking was performed in two steps: first with 3 % normal donkey serum and 0.3 M glycine in PBS for 30 minutes, and then with Mouse IgG Blocking Reagent (Vector Labs, Burlingame, CA). Primary antibody was applied (1:200, Mouse Monoclonal 235-1 IgG1, Rockland, Gilbertsville, PA) at room temperature for 1 hour, sections were rinsed with PBS, and the secondary antibody was applied for 20 minutes at room temperature (Alexa Fluor^®^ 488 Donkey anti-mouse, Invitrogen; 1:800). All nuclei were counterstained with DAPI (Vector Labs), and endogenous fluorescence was quenched with a 1 minute Trypan blue (250 μg/mL, pH 4.4, Invitrogen, Grand Island, NY) [21]. Sections were rinsed quickly with PBS, and mounted with aqueous medium.

### Quantification and analysis

To quantify the extent of vascular invasion into each group, 6 randomized images were taken from H&E-stained sections of each scaffold. The number of vessels, identified as lumina containing erythrocytes, was counted for each image, and 18 images (6 per scaffold; 3 randomized images per section at 2 depths) were averaged for each group and reported as the number of vessels per mm^2^ [22].

Cellularity was evaluated by quantifying the number of cells in DAPI-stained sections. Images were taken at central and peripheral radial locations of a given cross-section, at shallow and mid-height depths for each construct. The calculated number of cells from the center and periphery of a given cross-section were summed and normalized by the total imaged area to obtain a total number of cells per mm^2^. Images were taken at 400X magnification and processed in MATLAB (MathWorks, Natick, MA) with CellC (Selinummi & Seppala 2005; Wang et al. 2011), applying a segmentation factor of 0.9 and cell shape as the segmentation modality.

Statistical analyses were performed in Prism (GraphPad, La Jolla, CA). Significance was assessed by 1-way or 2-way ANOVA and Tukey’s post-hoc test (*p* < 0.05) as appropriate. Data are presented as the mean ± SEM.

## Results

### Cell seeding and scaffold preparation

Following the initial attachment period and overnight incubation, cell-containing constructs had a minimum seeding efficiency of 99.33 ± 0.11 % (Fig 1A). Histological analysis of scaffolds cultured for 14 days *in vitro* verified cellular infiltration throughout all scaffold types and a developing extracellular matrix in the pore spaces (Fig 1B-E). Cells appeared to be more evenly distributed in Col and CHA scaffolds compared to NuOss™ controls, where cells were more densely populated around the periphery of the scaffolds. NuOss™ (Fig 1B) and Col (Fig 1C) controls exhibited a non-circular cross-section and volumetric shrinkage after two weeks of culture, whereas CHA scaffolds cultured in either type of treatment medium (Fig 1D, 1E) maintained their original size and shape.

**Fig 1.**
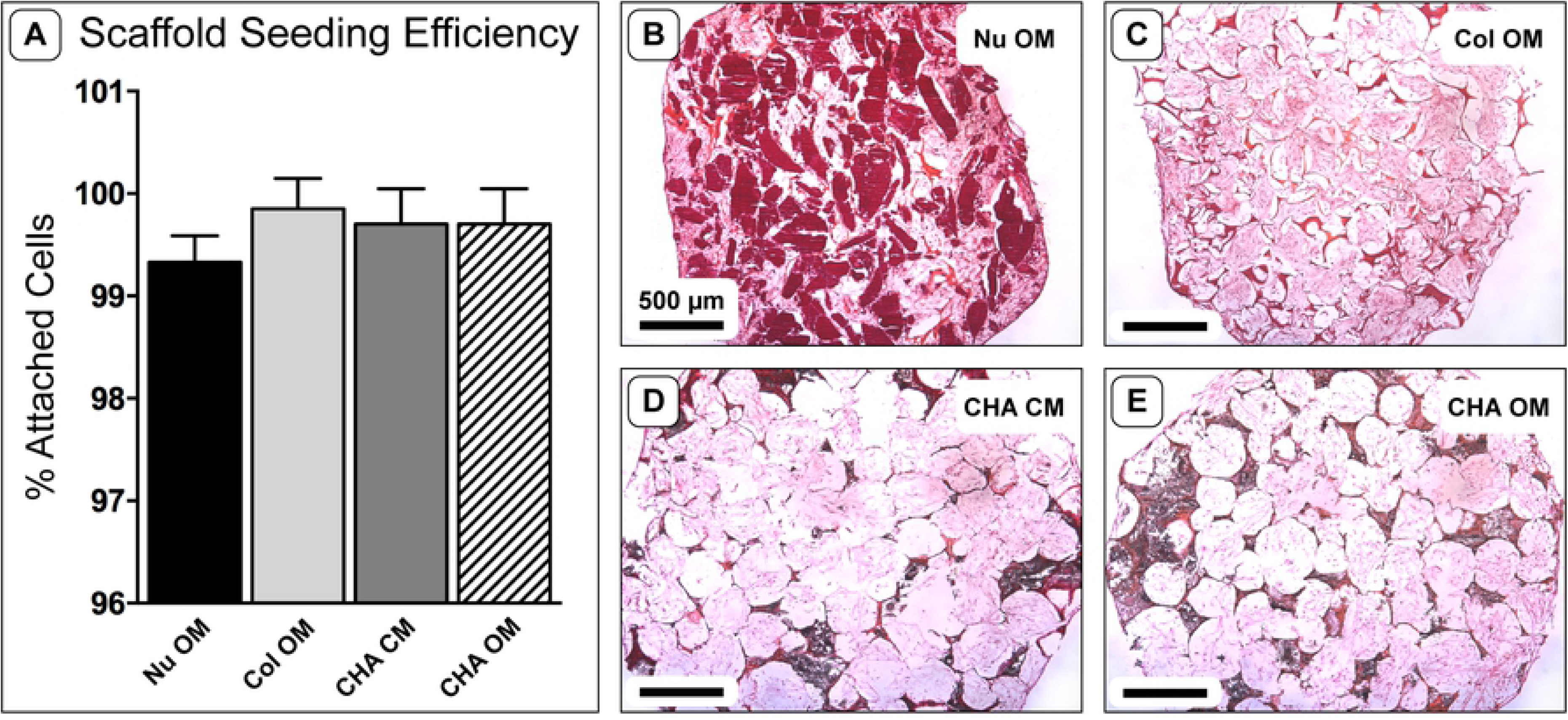
Scaffold seeding efficiency and morphology after *in vitro* culture. (A) Scaffold seeding efficiency, one-way ANOVA (*n* = 4; *p* = 0.17). (B-E) Representative H&E stained cryosections of hASC-seeded scaffolds after 14 days in culture. Note the composition of each scaffold by coloration: dark red indicates calcium phosphate granules in NuOss™ scaffolds; red-orange indicates collagen; black indicates the presence of HA whiskers; small, dark spots indicate cell nuclei; light pink indicates ECM deposited by hASCs; white space represents empty pore space.

Cells grown in monolayer and treated with OM and CM stained positive for alkaline phosphatase activity and contained small nodules of mineralization after 14 days of induction (not shown). The intensity and incidence of staining was higher for cells cultured with OM, indicating that this treatment was directing hASCs toward an osteogenic lineage.

### Tissue morphology and micro-CT analysis

Gross morphological evaluation revealed that Col and CHA CM explants were noticeably smaller than implanted constructs, and NuOss™ constructs were white, while all other groups had a light-pink tint (Fig 2). Blood vessels were most prominent on CHA OM and CHA Acel explants; however, vessels were also visible in CHA CM and Col explants. Interestingly, no blood vessels were visible in the NuOss™ control group.

**Fig 2.**
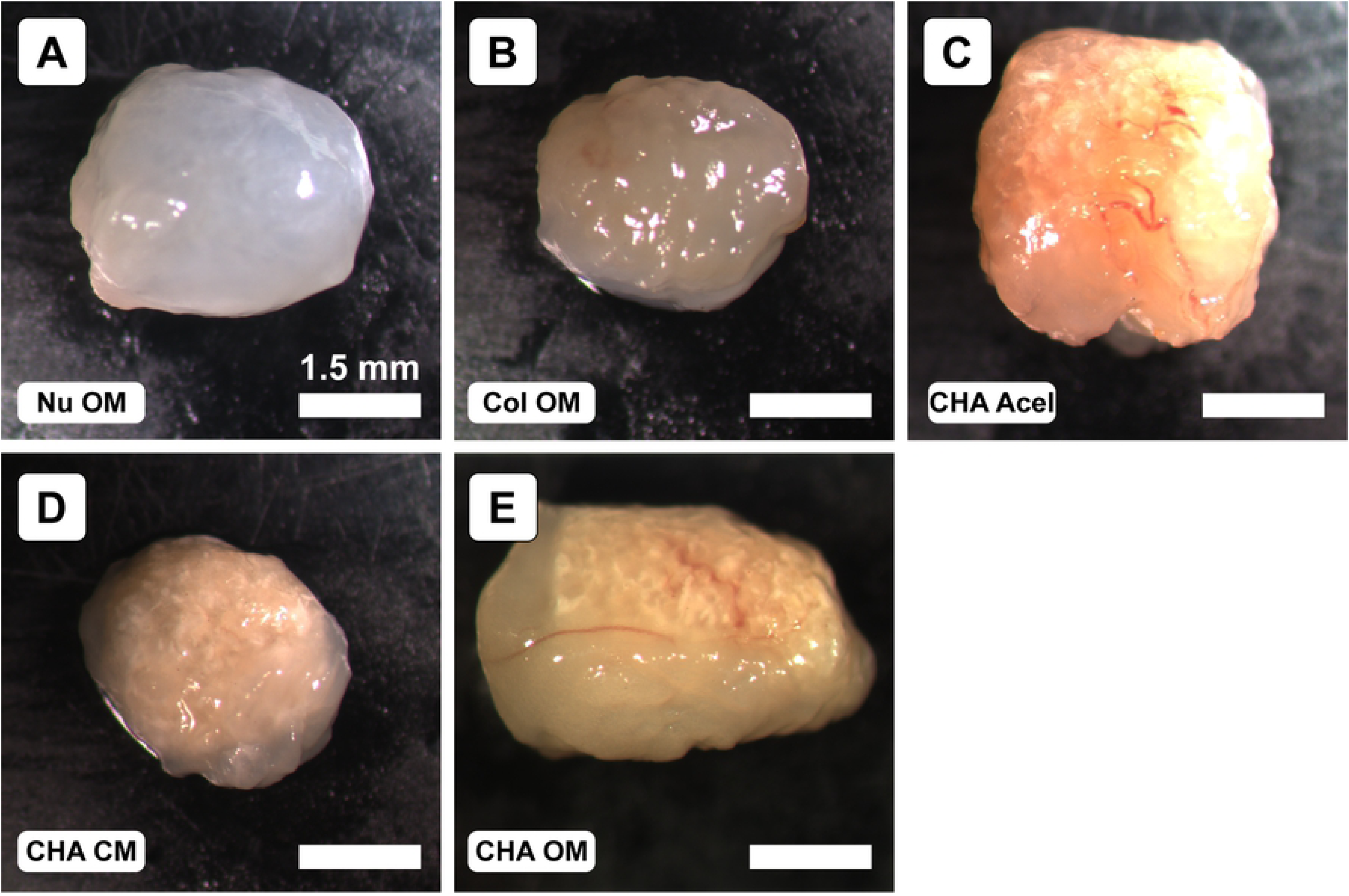
Representative gross morphology of explants after 8 weeks implantation.

After 8 weeks implantation, explants exhibited varying degrees of bone formation as determined by micro-CT. Each group exhibited a significant change between pre-implant (0 week) and post-explant (8 week) measurements of bone volume (Fig 3A). NuOss™ controls decreased in measured bone volume by 3.05 ± 0.55 mm^3^. In contrast, the bone volume of Col OM constructs increased from 0 to 1.06 mm^3^, and CHA OM constructs increased in volume by 2.08 ± 0.21 mm^3^. Interestingly, acellular CHA scaffolds increased in bone volume by 2.14 mm^3^, while the CHA CM constructs decreased by 2.40 ± 0.11 mm^3^.

**Fig 3.**
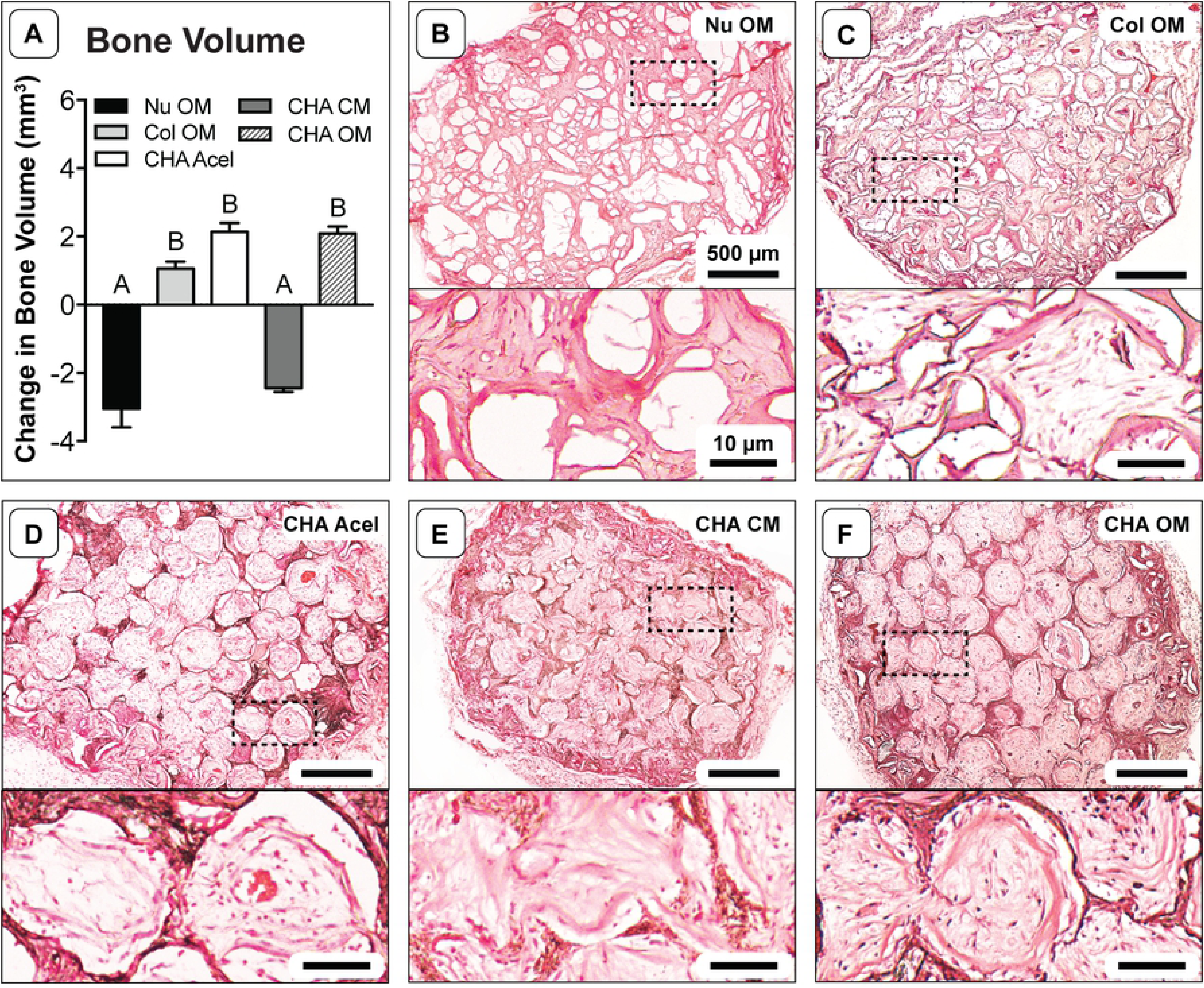
Change in bone volume and tissue morphology after 8 weeks implantation. (A) Absolute change in bone volume as measured by micro-CT; one-way ANOVA and Tukey’s post-hoc test (*n* = 3; *p* < 0.0001). Different letters indicate statistically significant differences. (B-F) Representative H&E stained cryosections showing the entire scaffold cross-section (upper image) and new tissue within scaffold pore spaces at higher magnification (lower image).

H&E stained sections revealed substantial differences in tissue morphology among groups. Like pre-implantation samples, the central region of NuOss™ scaffolds appeared to be less cellular than the periphery, based upon the eosin stain intensity (Fig 3B). Tissue infiltration in Col scaffolds (Fig 3C) was not as dense as in the CHA scaffolds (Fig 3D-F). CHA OM and CHA Acel scaffolds maintained a well-defined porous structure, while the pore structure of CHA CM and Col OM constructs appeared deformed and partially collapsed. Any changes in the porosity of NuOss™ scaffolds were less clear, but a certain amount of scaffold remodeling was apparent via tissue infiltration into the calcium phosphate granules.

Higher magnification revealed the organizational structure of the tissues within each group. Interestingly, treatment medium appeared to elicit a distinct tissue morphology and cellular response in hASC-seeded CHA scaffolds. CHA CM constructs formed a dense, disorganized tissue (Fig 3E); whereas CHA OM constructs produced a spatially organized tissue (Fig 3F). In general, the dense tissue observed in CHA OM constructs was preferentially located around the periphery of pore spaces and formed a ring-like structure around less dense tissue in the center of the pores. The tissue within CHA Acel constructs was similarly organized (Fig 3D). Dense, eosinophilic tissue was also observed at the edges of tissue within pore space in Col OM constructs; however, there were typically gaps between this tissue and the walls of the collagen struts (Fig 3C).

### Osteogenic and vasculogenic markers

Immunofluorescence staining revealed osteocalcin-positive tissue located in the matrix of CHA Acel and OM groups (Fig 4C,E). Positive staining was also observed to a lesser extent in NuOss™, Col and CHA CM groups (Fig 4A,B,D). Osteopontin immunofluorescence was more intense for HA-containing constructs, and appeared to localize to the scaffold structure (Fig 4H-J). Interestingly, the spatial patterning of osteopontin was similar in NuOss™ and Col groups, though the staining intensity was not as strong as in constructs that contained HA (Fig 4F,G).

**Fig 4.**
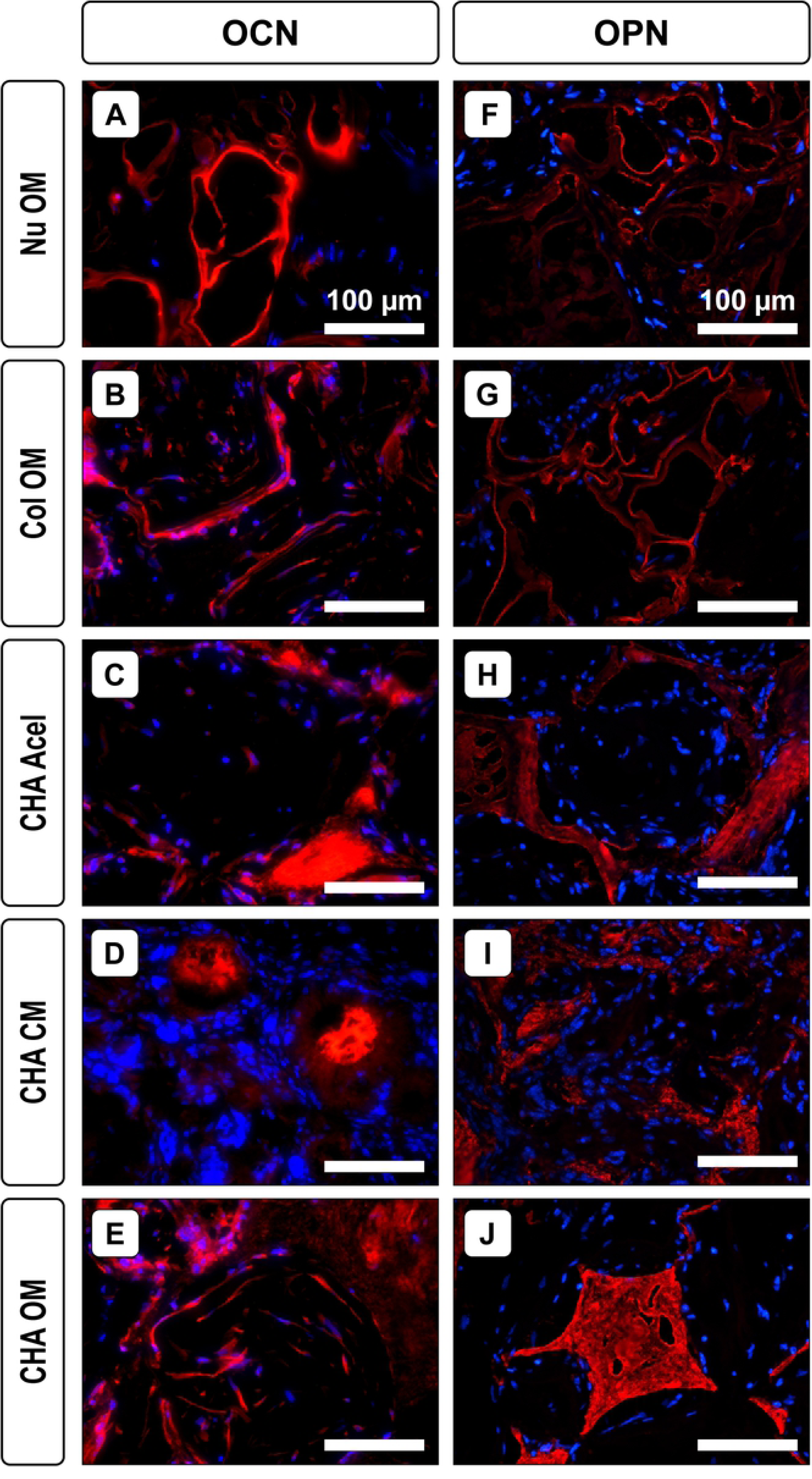
Osteogenic markers in explanted constructs. Representative immunostained sections showing (A-E) osteocalcin and (F-J) osteopontin after 8 weeks implantation. Red: osteocalcin or osteopontin; blue: DAPI (cell nuclei).

Histological analysis revealed the degree of vascularization for each group (Fig 5). CHA OM constructs had a higher blood vessel count (23.0 ± 3.3 vessels/mm^2^) than CHA CM (13.8 ± 2.1 vessels/mm^2^) and NuOss™ (5.0 ± 1.0 vessels/mm^2^) constructs. Col and CHA Acel also had significantly more vessels than NuOss™ scaffolds at 18.4 ± 1.8 and 16.8 ± 2.2 vessels per mm^2^, respectively.

**Fig 5.**
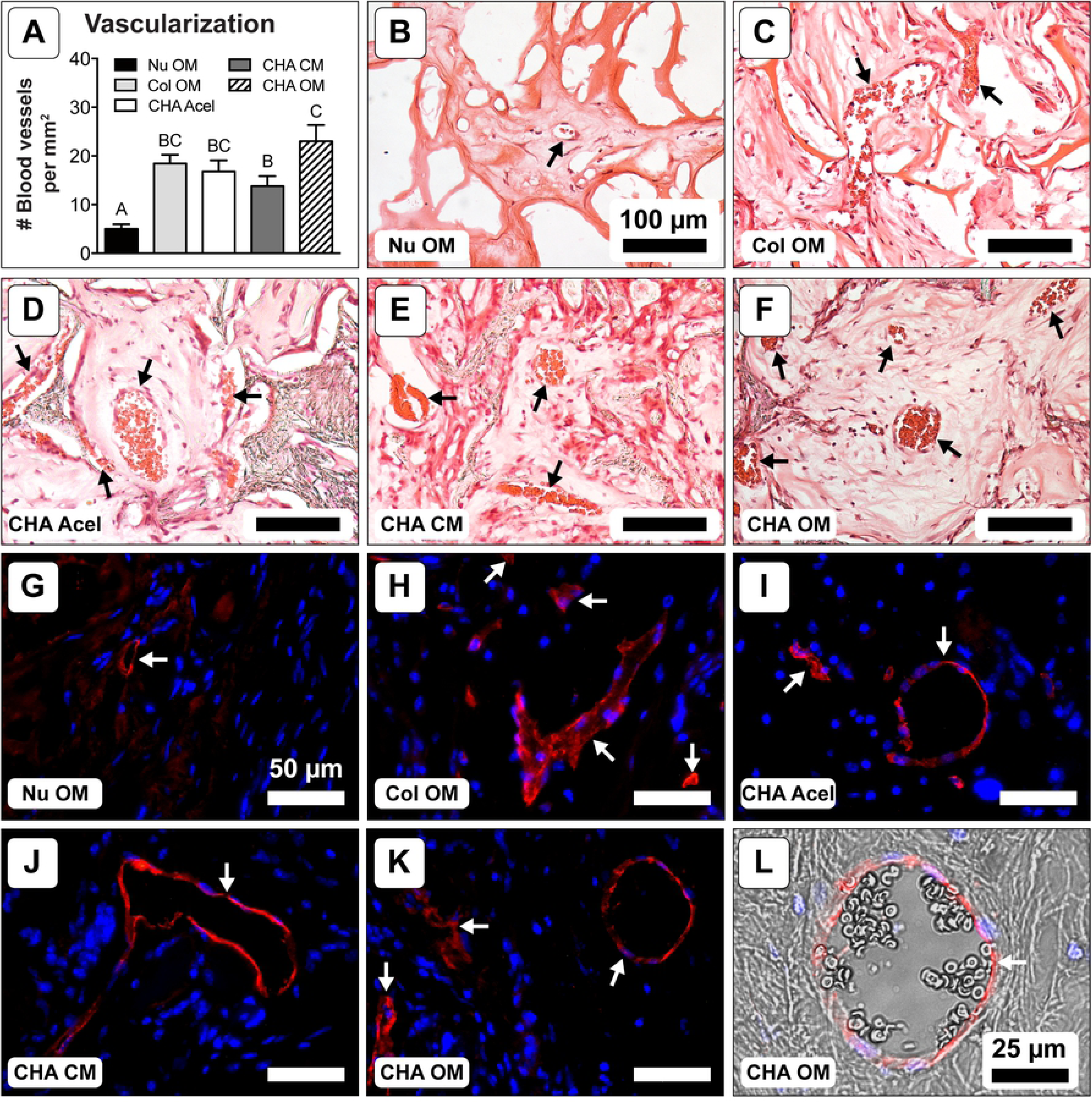
Vascularization after 8 weeks implantation. (A) Blood vessel density; one-way ANOVA and Tukey’s post-hoc test (*n* = 18; *p* < 0.05). Different letters indicate statistically significant differences. (B-F) Representative H&E stained cryosections showing blood vessels (black arrows). (G-K) Representative immunofluorescent stained sections showing CD31 (red) with DAPI counterstain (blue). White arrows denote blood vessels. (L) CD31 (red) and DAPI (blue) immunofluorescent staining overlaid on a matched bright field image showing the presence of erythrocytes within the vessel lumen.

To better understand differences in vascularization, VEGF levels in the cell-seeded scaffolds were also visualized via immunofluorescence. Within *in vitro* constructs, hASCs were actively secreting VEGF after 14 days of culture (Fig 6A-E, Pre-implant). Immunostaining for VEGF was more apparent in OM groups at this time point. Images of post-implantation constructs displayed a different trend (Fig 6F-J, Post-implant). CHA OM and CHA Acel groups had a high level of intense staining, while CHA CM constructs maintained a low level of VEGF. Nu OM constructs, which exhibited levels of VEGF staining comparable to CHA OM scaffolds after *in vitro* culture, exhibited a marked decrease in VEGF expression after 8 weeks of implantation. Staining was similar at both time points for the Col OM group; however, the intensity of the stain was lower than that of CHA Acel and CHA OM groups after 8 weeks of implantation.

**Fig 6.**
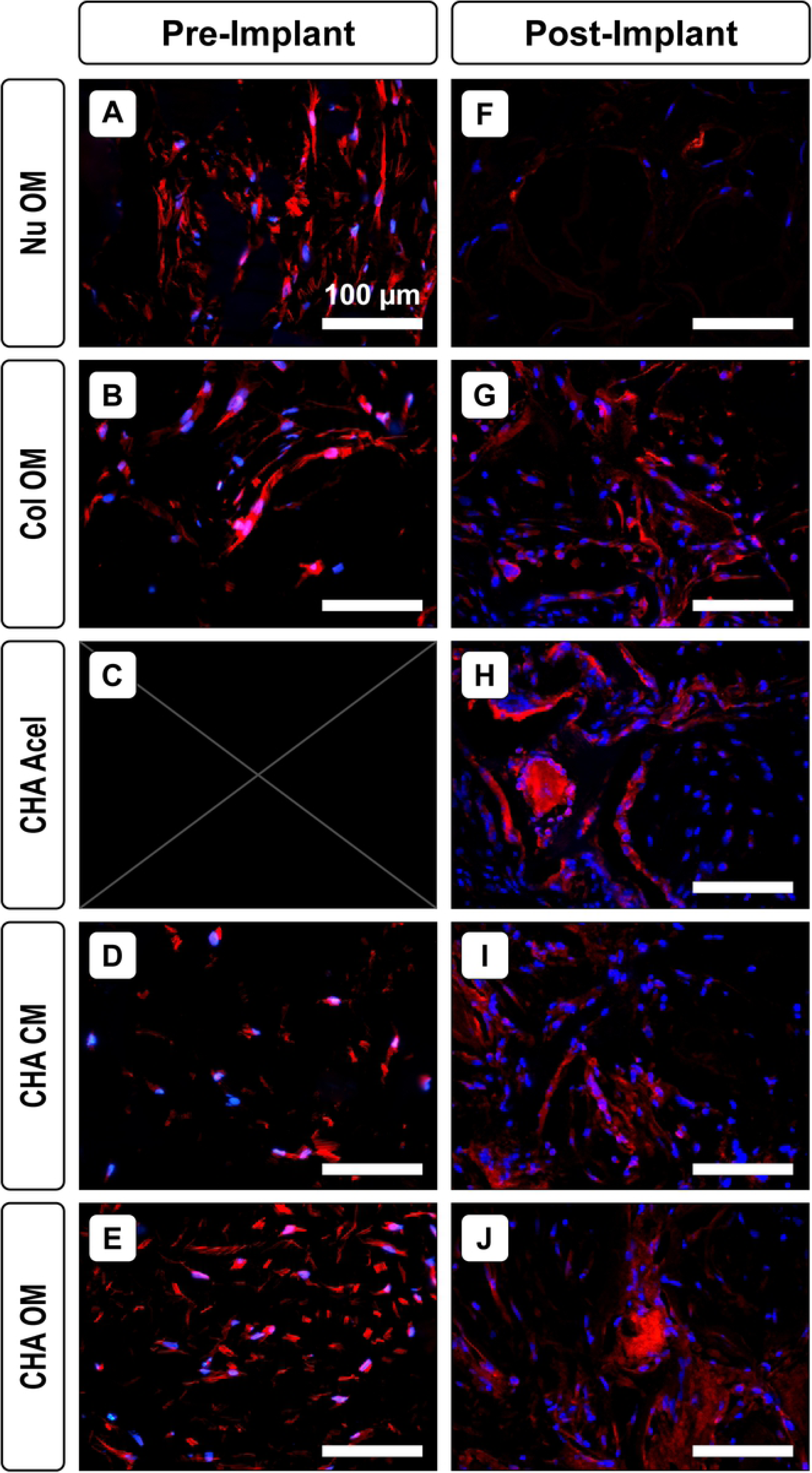
VEGF levels before and after implantation. Representative sections showing VEGF (red) and cell nuclei (blue) (A-E) after 14 days of *in vitro* culture but before implantation and (F-J) after 8 weeks implantation.

### Osteoclast activity and cellularity

Tartrate-resistant acid phosphatase (TRAP) staining demonstrated that there was considerable osteoclast activity in CHA CM constructs (Fig 7). Positive activity was also identified on the periphery of NuOss™ and, to a lesser extent, CHA OM explants. No staining was detected in Col or CHA Acel groups.

**Fig 7.**
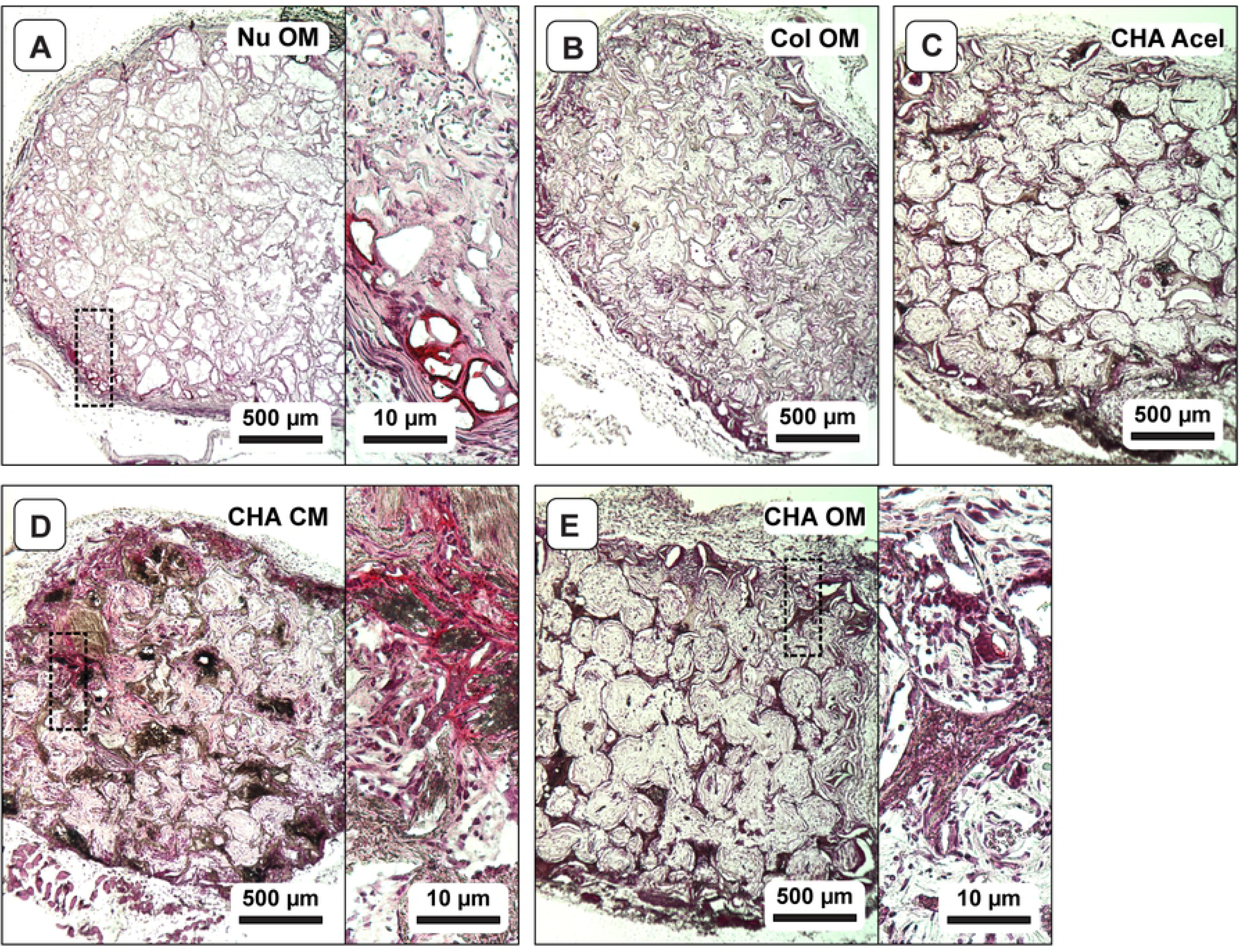
Osteoclast activity in explanted constructs. Representative TRAP-stained sections showing osteoclast activity (red).

The cellularity of each explant was also evaluated (Fig 8A). Comparing cell counts from the cross-sections sampled, the number of cells in constructs cultured *in vitro* for 14 days was not statistically different (Fig 8B). However, following 8 weeks *in vivo*, NuOss™ explants contained fewer cells and CHA CM explants contained more cells than all other groups (Fig 8B), with 8 week NuOss™ explants containing approximately the same number of cells as the 14 days *in vitro* time point. The cell distribution of evaluated cross-sections was not significantly different for Col OM and CHA CM constructs; however, the number of cells quantified at the center of CHA OM cross-sections was lower than at the periphery for *in vitro* scaffolds and for 8 week constructs (Fig 8B). Unlike NuOss™ explants, however, the cellularity of the CHA OM group increased by a factor of 4.5 following implantation.

**Fig 8.**
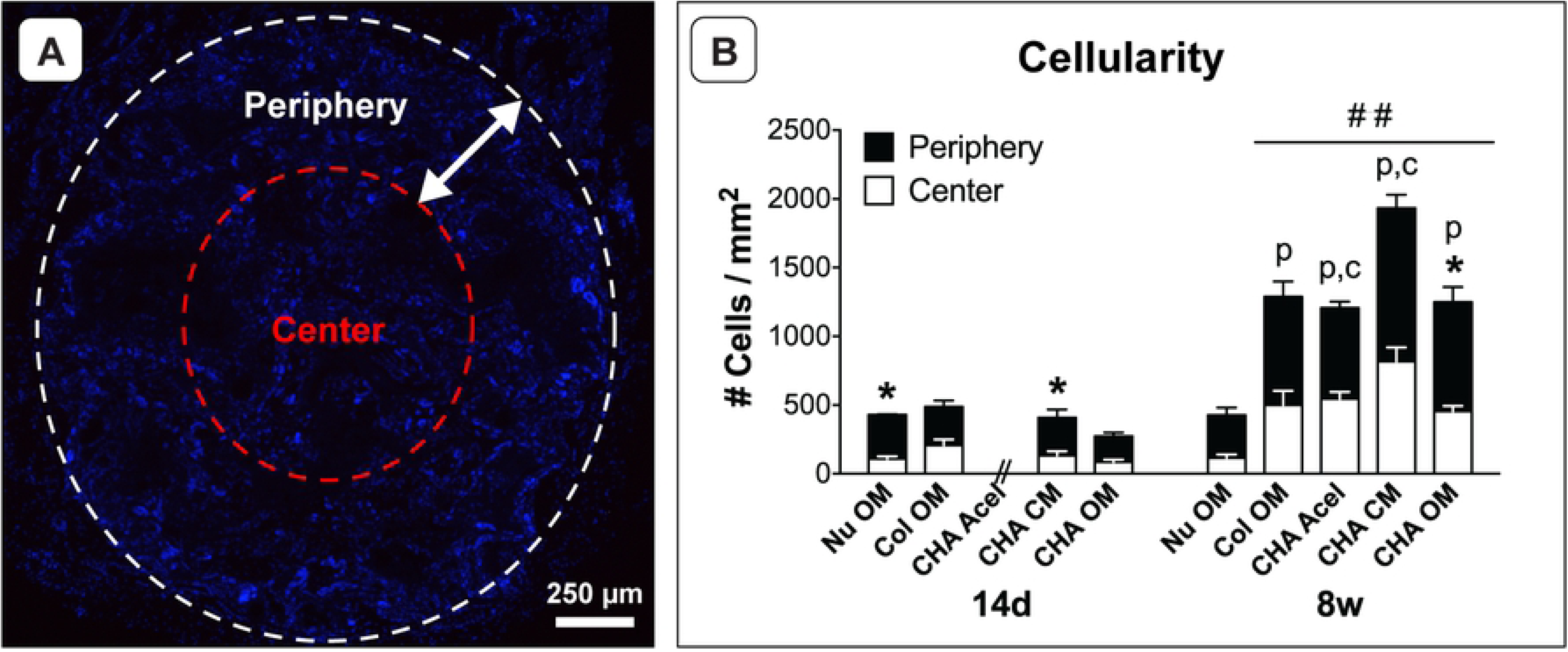
Cell distribution in scaffolds pre-and post-implantation. (A) Scaffold cross-section stained with DAPI. (B) Radial distributions of cells in *in vitro* samples (14d, *n* = 3) and 8 week explants (8w, *n* = 6). * indicates a significant difference between the periphery (black) and center (white) of individual scaffold groups (*p* < 0.05). ## indicates a significant increase in total cell number compared to 14d groups and 8w Nu OM (*p* < 0.001). Letters indicate that cell number for periphery (p) or center (c) significantly increased from 14d to 8w (*p* < 0.05).

Human nuclei immunofluorescence was used in conjunction with DAPI to evaluate the donor cell contribution to new tissue formation in explants. Human cells were identified within the implanted construct in all cell-seeded scaffold types (Fig 9); however, few remained after 8 weeks of implantation. The hASCs detected were primarily located on the periphery of the scaffolds or in the surrounding tissue. Additionally, despite physical separation within the mice, a single hASC was detected in the surrounding tissue of a CHA Acel explant (data not shown).

**Fig 9.**
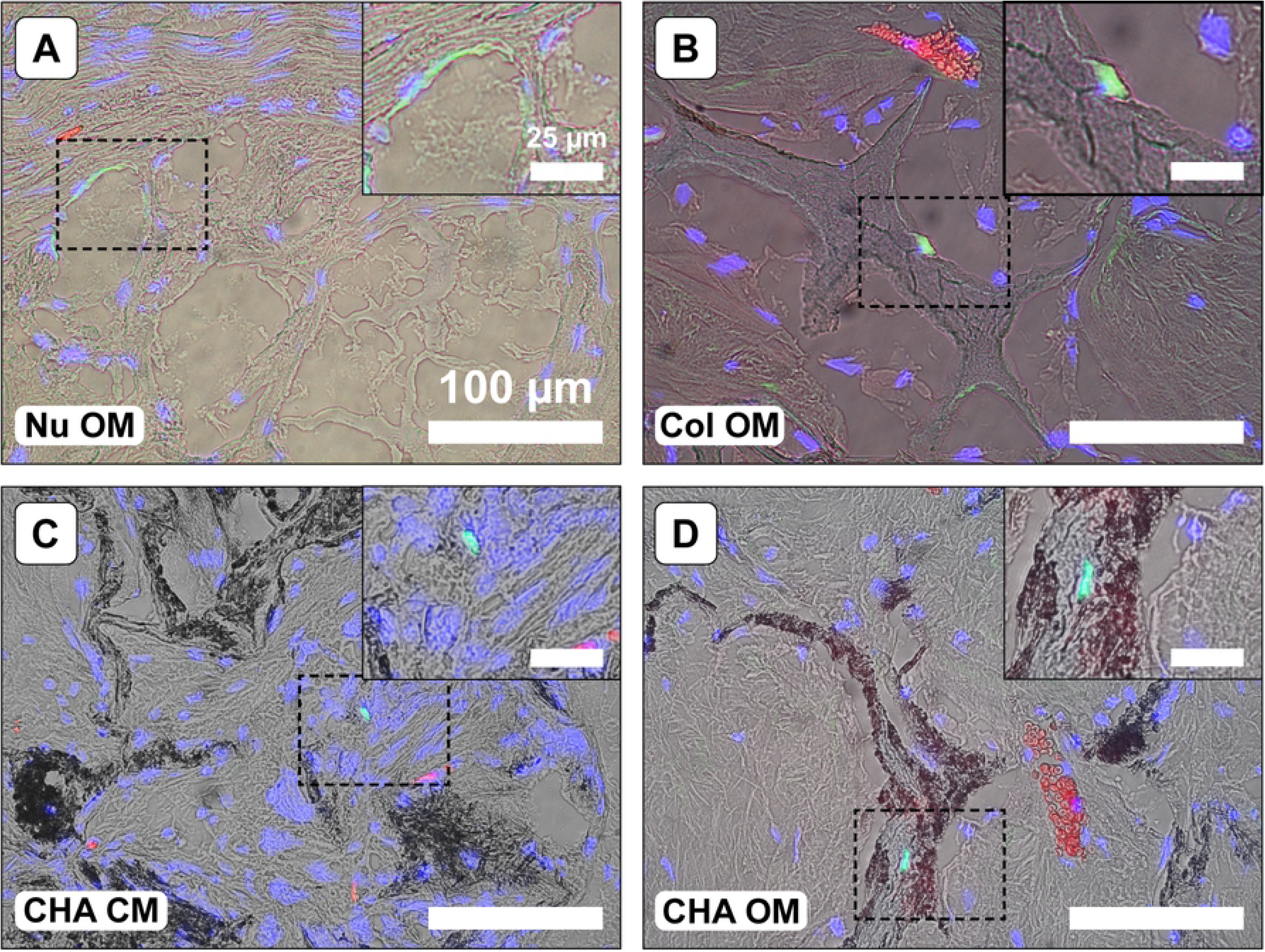
Detection of human cells within explanted constructs. Representative immunofluorescent stained sections showing human nuclei antigen (green) and cell nuclei counterstained with DAPI (blue) overlaid on a matched bright field image demonstrating the location of cells within the scaffolds. Blood vessels appear in red. The locations of magnified inset regions are indicated by black dotted lines.

## Discussion

The results of this study cumulatively indicate that treatment medium and scaffold composition direct osteogenic and angiogenic tissue formation in an ectopic model. One important effect of HA reinforcement in collagen-based carriers was evident after only 14 days of culture *in vitro*. All constructs with CHA scaffolds maintained the original size and circular cross-section throughout the pre-treatment period, whereas NuOss™ and collagen scaffolds exhibited volumetric shrinkage into an irregular, oblong-like shape. This finding suggests that HA whisker reinforcement increased the structural stability of collagen-based scaffolds throughout *in vitro* culture.

In terms of bone formation, the bone volume measured by micro-CT was increased in cell-seeded CHA scaffolds with pre-treatment in OM, but the increase was not significantly different from that in acellular CHA scaffolds. These results confirm previous reports that the CHA scaffolds alone promote the recruitment and osteogenic differentiation of endogenous cell populations [6,7]. The results further suggest that the choice of cell type and/or differentiation strategy should be carefully considered for pre-seeding these scaffolds. Additionally, ASC-seeded NuOss™ scaffolds exhibited in decreased in bone volume, suggesting that any bone formation was insufficient to counterbalance the resorption of the scaffold. Collectively, these results suggest that a combination of cell type and pre-treatment may need to be determined for a particular scaffold to achieve optimal bone regeneration. Similar results have been seen for other collagen-based scaffolds. Lyons et al. reported that matrix deposited by MSCs during in vitro culture may adversely affect healing by acting as a barrier to macrophage-mediated remodelling when implanted in vivo [25].

While pre-treatment of hASC-seeded CHA scaffolds in OM resulted in significant increases in bone volume, pre-treatment in CM resulted in decreased bone volume. This may be a result of the high osteoclastic activity observed via TRAP staining in the CHA CM scaffolds (Fig 7D). As opposed to mature osteoblasts, pre-osteoblasts have been reported to express higher levels of RANKL, which allows for the maturation, differentiation and activation of osteoclasts [26]; this may explain the high osteoclast activity in the scaffolds with undifferentiated cells in the current study. Osteoclast activity can be a positive indicator of scaffold resorption; however, the rate of resorption must be balanced with the deposition of replacement tissue and the maintenance of mechanical integrity.

To better characterize the tissue infiltrating the scaffolds, two bone markers were investigated: osteocalcin, a marker of osteoblasts that is associated with mineralized bone matrix; and osteopontin, a non-collagenous protein that is secreted by osteoblasts, osteocytes and osteoclasts and is therefore believed to play a role in both mineralization and bone remodeling [27]. Qualitatively, there was more osteopontin in CHA constructs than in collagen and NuOss™ scaffolds after 8 weeks of subcutaneous implantation. Localization of this protein to the scaffold structure is likely due to its ability to bind HA [27], and may account for the spatial organization of the tissue within CHA Acel and CHA OM groups. The majority of dense, bone-like matrix indicated by highly eosinophilic tissue (Fig 3) and concentrated osteocalcin staining (Fig 4) was identified at the periphery of the pore spaces in these constructs. HA may be at least partially responsible for this effect, as gaps were observed between the collagen-only scaffold and the extracellular matrix in the pores, and osteopontin levels were lower in these scaffolds.

Following implantation, the level of vascular invasion was higher in Col OM and CHA OM groups compared with CHA CM constructs, indicating that osteogenic pre-treatment may have contributed to this effect. Interestingly, CHA Acel controls achieved levels of vascularization comparable to CHA OM and Col OM constructs, while NuOss™ controls contained significantly fewer vessels. A similar trend was observed in VEGF levels: staining was stronger for CHA OM than for CHA CM, and there was considerably less VEGF detected in Nu OM constructs. VEGF, which is generally thought of as a key mediator of angiogenesis [28], also has the ability to regulate the recruitment and activity of osteoblasts, osteoclasts, and endothelial cells [29]. Therefore, higher levels of VEGF detected in CHA OM compared to CHA CM constructs both pre-and post-implantation may partially explain the differential bone formation and vascular invasion between these groups.

The tissue organization observed in CHA Acel and OM constructs was not maintained in CHA CM or Col OM tissues in the current study (Figs 1 and 3). For CHA CM scaffolds, this may be a result of the high osteoclastic activity observed via TRAP staining (Fig 7). The collapsed architecture apparent in the CHA CM group, along with a high level of osteoclast activity, were indicative of rapid resorption in these scaffolds. Decreased tissue organization in Col OM scaffolds could also be the result of collapsed pore structure, likely an effect of the absence of HA.

The decreased bone volume observed in NuOss™ scaffolds may be a result of its inhibited cellular infiltration compared to other groups. This is likely related to their 9% lower porosity compared to Col and CHA scaffolds, as reported by the manufacturer. The eosinophilic-rich, VEGF, osteocalcin and osteopontin-positive tissue in this group was observed only at the scaffold periphery, where the cell density was highest. This region coincided with the location of observed blood vessels and osteoclast activity. Conversely, the central region had a very low cell density both pre-and post-implantation, and bone markers were not detected in this area. Previous studies with hASCs have reported that cell density has a significant impact upon resulting tissue formation [30], indicating that a lower concentration of cells at the scaffold center may have influenced its bone forming capacity.

Despite significant differences in vascularity, bone formation and cellularity in cell-seeded groups, few human cells were identified in any of the explants. Remaining hASCs were primarily located at the periphery of the scaffold, or in the surrounding tissue, perhaps indicating their migration out of the constructs. This hypothesis is supported by the detection of a human cell in the tissue surrounding an acellular construct. Both retention [30] and loss [31] of ASCs has been reported in the literature, introducing further confusion regarding their role in ectopic bone formation.

## Conclusions

The results of this study indicate that both scaffold type and pre-treatment are crucial to successful mineral deposition and vascular invasion, and that to achieve optimal bone formation it may be necessary to match the scaffold with a particular cell type and cell-specific pretreatment. HA-reinforcement allowed collagen constructs to maintain their implanted shape, provided for improved cell-tissue-scaffold integration, and resulted in a more organized tissue when pre-treated in an osteogenic induction medium.

